# DDIA: data dependent-independent acquisition proteomics - DDA and DIA in a single LC-MS/MS run

**DOI:** 10.1101/802231

**Authors:** Shenheng Guan, Paul P. Taylor, Ziwei Han, Michael F. Moran, Bin Ma

## Abstract

Data dependent acquisition (DDA) and data independent acquisition (DIA) are traditionally separate experimental paradigms in bottom-up proteomics. In this work, we developed a strategy combining the two experimental methods into a single LC-MS/MS run. We call the novel strategy, data dependent-independent acquisition proteomics, or DDIA for short. Peptides identified by conventional and robust DDA identification workflow provide useful information for interrogation of DIA scans. Deep learning based LC-MS/MS property prediction tools, developed previously can be used repeatedly to produce spectral libraries facilitating DIA scan extraction. A complete DDIA data processing pipeline, including modules for iRT vs RT calibration curve generation, DIA extraction classifier training, FDR control has been developed. A key advantage of the DDIA method is that it requires minimal information for processing its data.

**GRAPHIC ABSTRACT:** 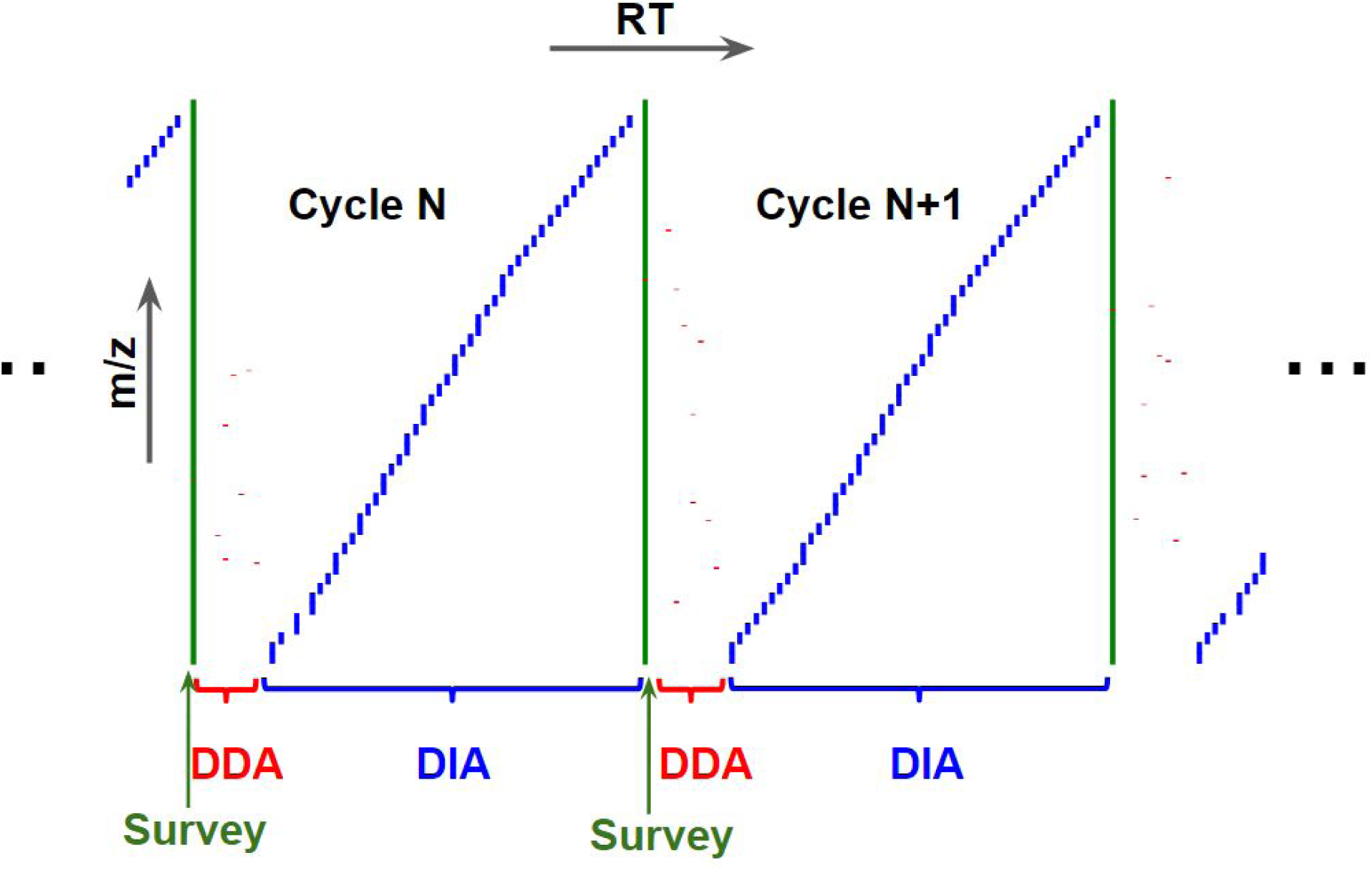

## INTRODUCTION

High performance mass spectrometry based bottom-up proteomics offers in-depth analysis of proteins in samples of complex mixtures, by identifying and quantifying corresponding peptides. Two general classes of bottom-up or shotgun proteomics experiments are data dependent acquisition (DDA) and data independent acquisition (DIA) methods.

The DDA (data dependent acquisition) methods have been extensively practiced in the proteomics field since its inception. One widely used implementation of DDA experiments is the top N method, in which the N most abundant precursor ions were selected for fragmentation. Dynamic exclusion and other techniques allow for achieving deeper precursor coverage and avoiding repeated acquisition of most abundant precursor ions. Peptide identification in DDA experiments is generally realized by protein sequence database search of fragment spectra. Target-decoy sequence database search can be used effectively for false discovery rate control of DDA peptide identification.

In DIA (data independent acquisition) experiments, a set of predetermined wide (often overlapping) isolation windows is systematically used to send precursor ions in an isolation window for fragmentation. With sufficiently wide isolation window coverage, all precursor ions are subjected for analysis.

Due to multiplexing nature of DIA ms2 spectra, DIA data analysis is challenging. The robust protein sequence database search methods and target/decoy strategies in DDA experiments cannot be applied directly. Several DIA data processing algorithms have been developed. MSPLIT-DIA(1) uses spectral similarity to extract DIA data. DIA-Umpire(2) extracts pseudo spectra from DIA data and identifies the pseudo spectra with DDA-like protein sequence database search engines. Group-DIA(3) generates pseudo spectra by deconvolution of precursor-fragment pairs from several DIA data files. The most widely used DIA data analysis methods are still the use of a spectral library constructed from DDA experiments to extract ion chromatograms from DIA data. OpenSWTH(4), Spectronaut(5), and Skyline(6) are a few widely used software packages. For in-depth interrogation of a DIA experiment, the spectral library needs to be sufficiently large and retention times of peptide ions in spectral library should correlate well with the DIA data of interest. To address these issues, we have developed deep learning based tools(7) to predict LC-MS/MS properties from peptide sequence for DIA data analysis. LC-MS/MS properties of peptides, such as indexed retention times (iRT), ms1 charge state distributions, and HCD spectra can be predicted accurately from their sequences.

Leveraging on the capabilities of our LC-MS/MS property prediction tools, we introduce here DDIA: a data dependent-independent acquisition method, a new MS instrument method combined with a novel data analysis strategy. In a DDIA experiment, both DDA and DIA are combined in a single LC-MS/MS run. A DDIA cycle of scans includes a survey scan followed by both DDA ms2 scans and DIA ms2 scans. A major portion of cycle time is spent on DIA scans and a small number of high quality DDA scans can be obtained with a small portion of cycle time. The DDA scans can be identified by conventional sequence database search. Identification of peptides in the DDA scans provides useful information for interrogation of the DIA scans. The following information can be obtained from analysis of DDA identified peptide ions: (a) an indexed retention time (iRT) vs retention time (RT) calibration curve, and (b) a classifier for DIA extraction FDR control. A larger spectral library, for example for high detectability peptides(8) in human proteome, can then be constructed by spectral prediction tools for in depth interrogation of the DIA scans.

We demonstrated the DDIA strategy with analysis of a HeLa lysate digest sample. The DDIA mass spectrometry method has a cycle time of 3.6 seconds, in which 0.6 second is on a survey scan and DDA scans and the rest of 3.0 seconds on DIA scans. A complete data processing pipeline was built to process DDIA data from raw files to DIA extraction identification with false discovery control. The LC-MS/MS property prediction tools(7) were repeatedly used in the pipeline to generate spectral libraries for DIA extraction. False discovery rate (FDR) control was realized using a decoy spectral library created from a decoy sequence database.

The core advantage of the DDIA method is that minimal amount of information needs to be fed into the data processing pipeline. Indeed, the only assumption made for the sample is its species of origin. Therefore, the DDA identification and the high detectability peptide list generation can use the same proteome sequences of the species. There is no need to spike in iRT standard peptides into samples for retention time calibration and to prepare experimental spectral libraries from separated DDA experiments.

## EXPERIMENTAL PROCEDURES

1.5 **μ**g of Pierce HeLa Protein Digest Standard (PN:88328,Waltham, Massachusetts) for each run was used.

The DDIA experiment was carried out on an Orbitrap Fusion™ Tribrid™mass spectrometer from ThermoFisher Scientific (San Jose, California) interfaced with an Easy nLC 1000 nanoflow liquid chromatography (also from ThermoFisher Scientific). The peptides were separated on a 50 cm long Easy-Spray Acclaim PepMap column with 100Å pore size and 75µm ID. The nLC was running at 250nL/min and with a gradient program of 1) 5%B at 0 min to 34%B at 106 min; 2) 90%B for 14 mins. Solvent A was 0.1% formic acid in water and solvent B was 0.1% formic acid in acetonitrile. The Easy-Spray column was kept at 60°C.

The MS instrument method was constructed with two concurrent cycle experiments. The experiment 1 with cycle time of 0.6sec consists of a ms1 or survey scan and data dependent (DDA) scans. The survey scans were acquired with a resolution of 60k with a scan range of 400 - 1600. The The DDA scans were acquired in the orbitrap with 15k resolution, 30% HCD collision energy, and an isolation window width of 1.6. The experiment 2 with 3sec cycle time has 55 data independent (DIA) scans, with an isolation widow width of 12 and window center spacing of 10. So there is 1 m/z unit overlapping at the boundary of two adjacent scans. The DIA scans were acquired at 15k resolution, scan range 250 - 2000, and 30% HCD collision energy. The lowest and the highest DIA scan centers are 455.4775 and 995.7475, respectively.

### DDIA data processing pipeline

The DDIA raw data files were processed using home-developed data processing pipeline, which incorporates several published software modules. The major portion of the pipeline was written in Python. It’s architecture and major modules are described in the Results section and supplemental materials. Five components of the pipeline were published modules described below.

#### Data access

The survey scans, DDA ms2 scans, DIA ms2 scans, and the instrument method were extracted from the raw data files and saved separately using previously developed PAVA code(9).

#### De Bruijn decoy protein sequence generation

Decoy protein sequences were generated from normal protein databases using the de Bruijn decoy generation tool(10), which creates a decoy protein sequence from a target protein sequence, preserving repeating peptides in a normal (target) sequence database.

#### Sequence database search of DDA ms2 scans

The sequences of human proteome (Uniprot Proteome ID: UP000005640 with 71778 entries) were downloaded on February 6, 2018. The corresponding de Bruijn decoy sequences were concatenated by use of the target sequences. The DDA ms2 peaklists were searched with the MSGF+ search engine(11) against the concatenated sequence database. Trypsin specificity with up to 5 missed cleavages were allowed (filtered after search). Cysteine carbamidomethylation was the fixed modification. Variable modifications were oxidation of methionine, pyro-glu from peptide n-terminal glutamine, and protein n-terminal acetylation with or without loss of methionine, and deamidation of glutamine and asparagine. Precursor m/z tolerance was 10ppm and fragment ion detection instrument was set to “Q Exactive”. The peptide EValue cutoff of 0.65 was used to obtain the peptide ion FDR of 1.0%.

#### Peptide LC-MS/MS property prediction tools

LC-MS/MS properties, indexed retention time (iRT), ms1 or survey scan charge state distribution, and HCD spectral intensities of peptides of interest were predicted from their sequences (and precursor charge state for HCD spectral prediction) using deep learning models(7). For prediction performance comparison, predicted spectral libraries of peptides of interest were also obtained from the Prosit web service(12).

#### Peptide detectability prediction

Peptide detectabilities were computed by the AP3 program(8).

## RESULTS

Although the instrument method and the data processing algorithms have not been extensively optimized, the data dependent-independent acquisition (DDIA) experiment clearly demonstrated its capability: significant numbers of protein groups can be identified with minimal assumptions about the sample. The only assumption was the sample of human species. The data processing pipeline was able to learn all necessary information, from the raw data file.

### DDIA instrument method

As illustrated in Figure 1, the DDIA instrument method consists of a survey or ms1 scan followed by data dependent (DDA) scans, and then by data independent (DIA) scans. The cycle time of the survey scan and the DDA scans was set to 0.6 seconds. The cycle time for the 55 DIA scans was 3.0 seconds. Therefore, it takes 3.6 seconds to complete one whole cycle and the instrument spent about 2% of time on survey scans, 15% on DDA scans, and 83% on DIA scans.

**Figure 1.**
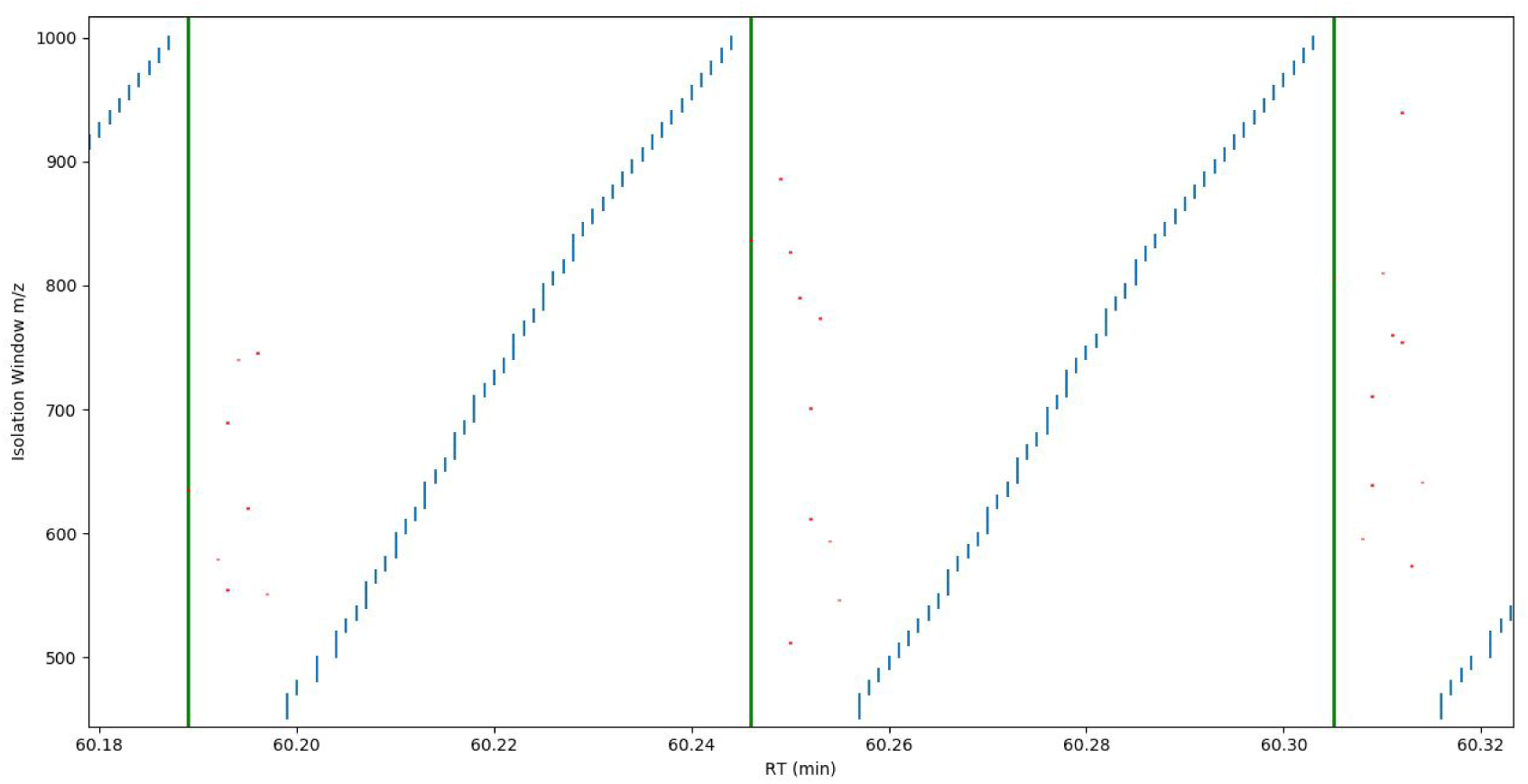
A portion of retention time vs ms1 m/z space targeted by the DDIA method. Two complete cycles of scans are shown. Each vertical line segment indicates the retention time (the horizontal axis) and the length of scan range (survey) or ms2 isolation window width of a scan. The green, red, and blue vertical line segments are for survey, DDA, and DIA scans, respectively.

In the present method, ions in all scans were detected in Orbitrap. High mass resolution and accuracy of the Orbitrap detection are useful for survey and DIA scans. Detection of DDA scans in Orbitrap was only useful to allow for spectral similarity comparison between DDA spectra and DIA pseudo spectra and between spectral prediction models in this study. A more optimized DDIA method should have DDA scans be detected in ion trap to provide higher sensitivity and speed.

### DDIA Data Processing Pipeline

The architecture of data processing pipeline is shown in Figure 2 and the central scheme is the backbone of four processing modules of (a) the previously published LC-MS/MS property prediction tools(7), (b) spectral library construction module, (c) DIA extraction module, and (d) DIA feature classifier.

**Figure 2.**
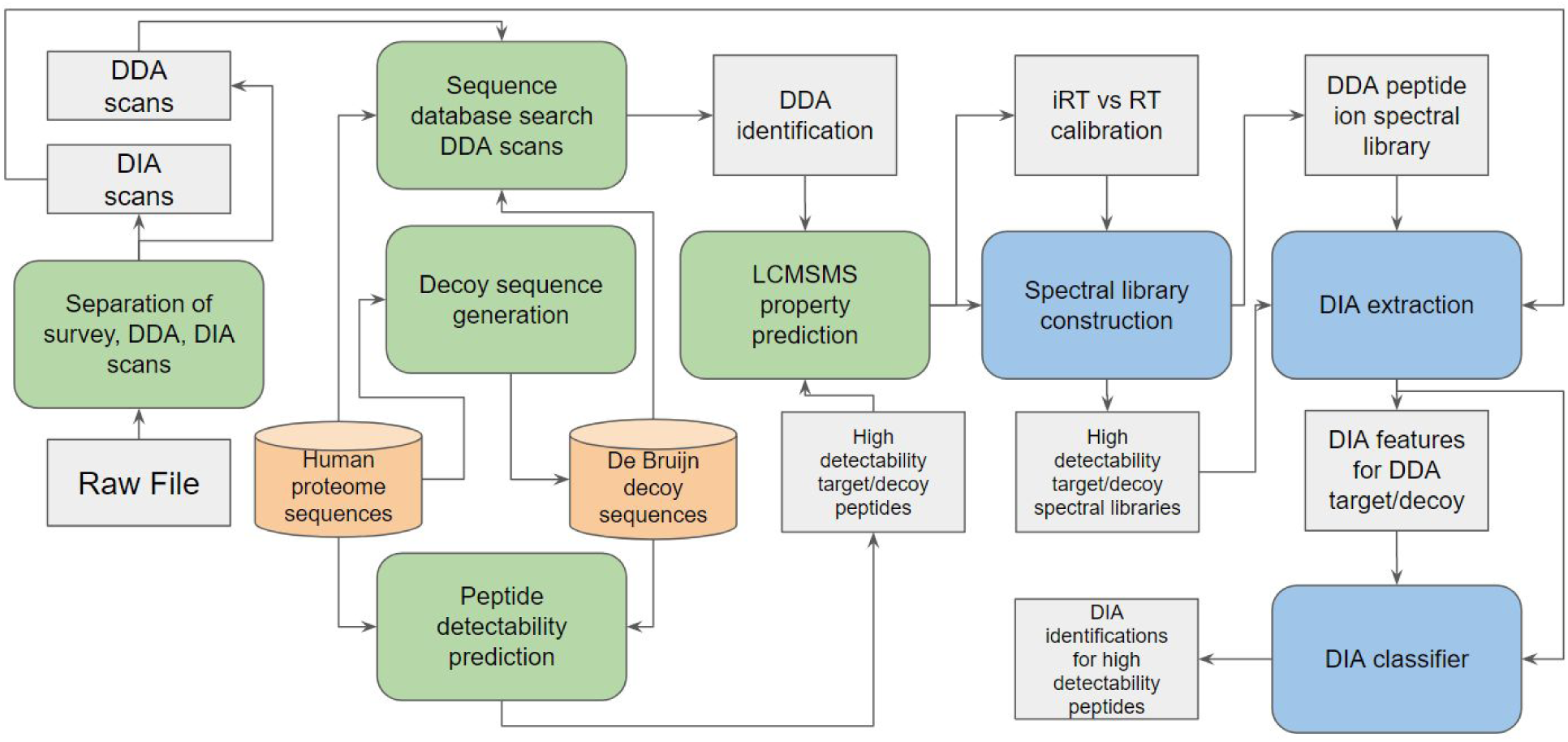
Flow chart of DDIA data processing pipeline. Rectangular shapes are data blocks and round corner blocks are processing modules. The light green shaded blocks are previously published processing tools.

The information flows through the DDIA data processing pipeline in four logical stages.

Processing stage 1 (supplemental figure S1): The iRT vs RT calibration curve and target DIA features for DDA identified peptides were obtained. The target spectral library was constructed using predicted HCD spectra. Retention time for each of the spectral library entry was calibrated through its predicted iRT value via the iRT vs RT calibration curve. The target DIA features were extracted from the DIA scans and they would be used to train a DIA extraction classifier.

Processing stage 2 (supplemental figure S2): The decoy DIA features were obtained by extraction from DIA scans with downsampled (matching the number of the target peptides) high detectability peptides of the decoy protein sequences (de Bruijn decoy).

Processing stage 3 (supplemental figure S3): The DIA extraction classifier was trained using the target DIA features for DDA identified peptides (output of Processing stage 1) and the decoy features from the de Bruijn peptides (output of Processing stage 2).

Processing stage 4 supplemental figure S4): The DIA features for high detectability peptides were extracted and classified by the DIA extraction classifier to obtain positive identifications.

### Identification of DDA peptide ions

A DDIA raw data file of two hour run was chosen to illustrate the processing pipeline.

The workflow starts with separation of the raw data file into the instrument method, survey scans (1870), DDA scans (15336), and DIA scans (102850). The DDA ms2 scans were subjected to sequence database search against the human proteome sequences and the corresponding de Bruijn decoy sequences(10). 8217 peptide ions were identified with the false discovery rate (FDR) of 1%.

### Construction of iRT vs RT calibration curve

Accurate retention times (RT) of the DDA identified peptides can be obtained by extraction of precursor chromatograms. For example, the isotope ion chromatograms of the precursor ion may be extracted from survey scans and the XIC was fitted with a polynomial variance Gaussian function(9). But this seems not necessary, since the median chromatographic peak widths of 0.255 mins is less than an order of magnitude of the retention time extraction window width determined by iRT prediction accuracy. The retention times of the identified DDA scans were directly used instead.

Indexed retention times (iRT) of the DDA identified peptides were predicted from their sequence using the LC-MS/MS property prediction tools(7). The iRT vs RT calibration curve was constructed using a moving median method with a window width of 4 iRT unit. Figure 3 is a scatter plot of iRT vs RT with the calibration curve and ±3 min boundary bands.

**Figure 3.**
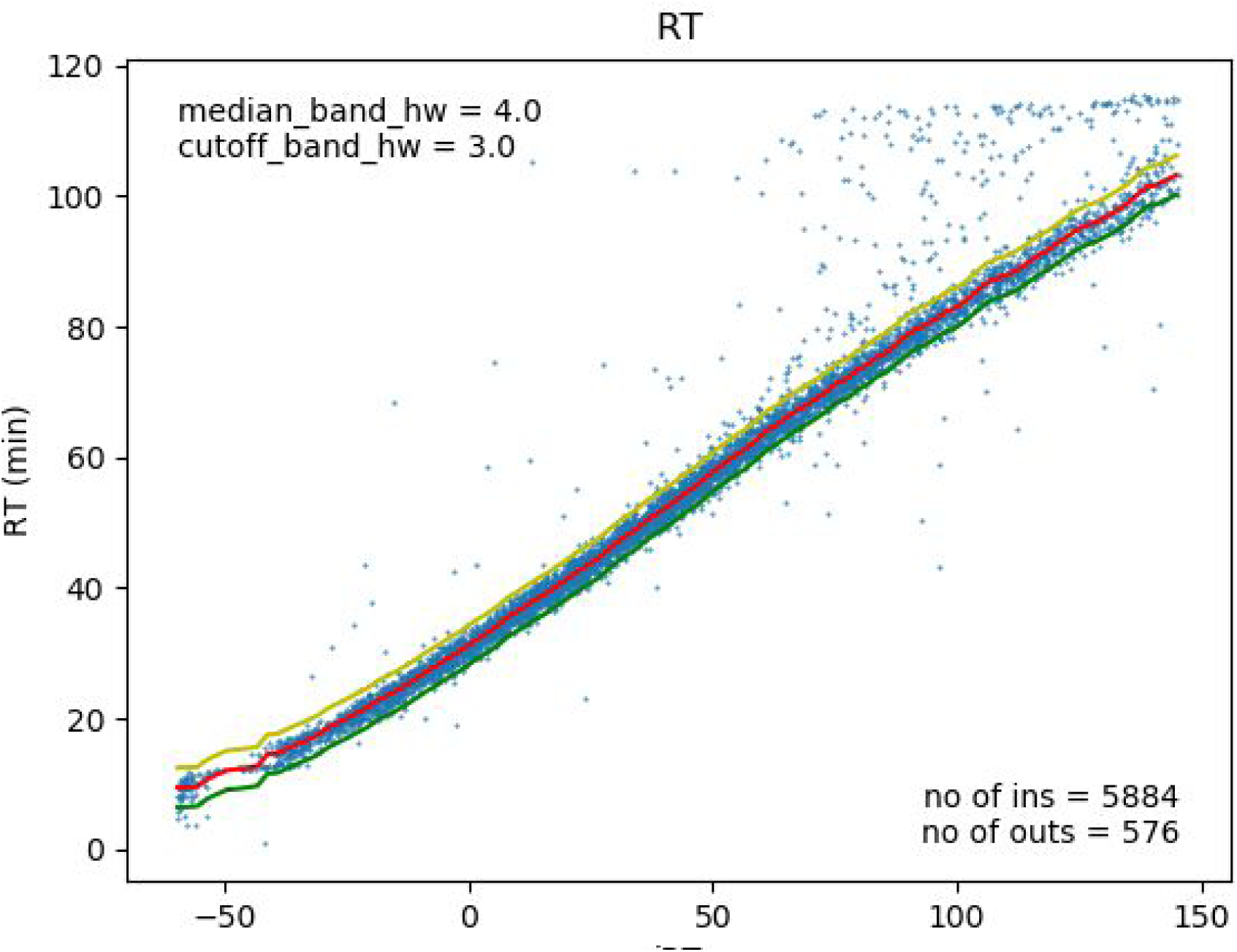
Scatter plot of iRT vs RT for the DDA identified peptides. The calibration curve (red) was obtained using a moving median method with window width of ±4 iRT unit. Yellow line is the upper boundary band(+3 RT min from the calibration curve). Green line is the lower boundary band(−3 RT min from the calibration curve).

There are many points in the plot are higher than the median calibration curve, especially in the upper right corner. Those peptides may reside on the column during the normal gradient and may be washed out in the final rapid ramp of the gradient. The data aggregation using moving median allows for rejection of those outliers. The accuracy of iRT prediction can be estimated by the tightness of iRT vs RT points around the calibration curve.

### Construction of three spectral libraries

#### Target spectral library of DDA identified peptide ions for training DIA extraction classifier

For each of the 5884 DDA identified peptides, which reside in the ±3 mins boundary bands of the iRT vs RT calibration curve, the preferred targeted charge states were determined by the predicted charge state distribution by the LC-MS/MS property prediction tools(7), if the precursor m/z is in the range of DIA scans (455.4775 - 5 to 995.7475 + 5). The resultant 8124 peptide ions were filtered (remove modified peptides) and the filtered list of peptide ions were submitted for HCD spectral prediction(7). The predicted HCD spectra and their retention time obtained from the iRT vs RT calibration curve were combined into the target spectral library (7781) for DDA identified peptides. Ten most abundant ions were selected.

#### Decoy spectral library for training DIA extraction classifier

Human proteome sequences (71,778 protein entries) were converted to de Bruijn decoy sequences(10). The AP3 program(8) was used to provide a list of peptides with their detectability from the de Bruijn decoy database. 568,117 peptides were selected with the detectability greater and equal than 0.6. As discussed in the section for DDA identified spectral library construction, the peptide’s charge state can be selected in the range of DIA scans. HCD spectra were predicted for the 704,063 peptide ions and their retention times were obtained from their predicted iRT values using the iRT vs RT calibration curve. 7781 entries of the de Bruijn decoy spectral library were randomly selected as the decoy spectral library for training the DIA extraction classifier.

#### Construction of High Detectability Peptide Spectral Library

High detectability target peptide spectral library was constructed similarly as for the de Bruijn decoy spectral library, except this time the human proteome sequences were directly used. There are 551,628 peptides with the detectability greater and equal than 0.6 in the human proteome sequences according to the AP3 program(8). The final high detectability peptide spectral library has 704,609 entries.

### DIA Extraction

Ion chromatograms from survey scans and DIA scans were extracted using the spectral libraries described early. For a precursor, its isotope distribution was calculated and isotope fine structures were centroided to give mass and intensity pair for each nominal mass above an intensity threshold of 20% of maximum. After adjusting for its charge state, isotope ion chromatograms from survey scans were collected. Fragment ion chromatograms were extracted for upto 10 fragment ions in a spectral library entry from DIA scans with the corresponding isolation window. The m/z tolerances were 10ppm and 25ppm for survey and ms2 extractions, respectively. The retention time extraction window was ±3mins determined by the iRT vs Rt calibration plot (figure 3).

Extraction and aggregation of DIA chromatograms is described in detail in Supplemental Material. Briefly, the extraction module extracts isotope ion chromatograms from survey scans and aggregate them into dot product and pairwise distance (against theoretical isotope distribution) chromatograms and extract fragment ion chromatograms from DIA ms2 scans and aggregate them also into dot product and pairwise distance (against spectral library product ion intensities) chromatograms. The four chromatograms were then aggregated together into a peptide ion chromatogram, from which the peak duration was determined and 6 features for the peptide ion were computed within the peak duration.

### DIA Extraction Classifier Training

7781 feature vectors for the DDA identified target spectral library and 7781 feature vectors for the downsampled de Bruijn decoy spectral library were given label of class 1 and 0, respectively. The 13221 vectorts of features were randomly split into the training set (11347) and testing set (3783). The training set was used to train a Quadratic Discriminant Analysis (QDA) classifier. For the testing set, the false discovery rate of 1.06% was achieved with the classification probability threshold of 0.996.

The performance of the DIA extraction classifier is shown in the confusion matrix format in Table 1 and the receiver operating characteristic (ROC) curve in supplemental figure S5.

**Table 1.**
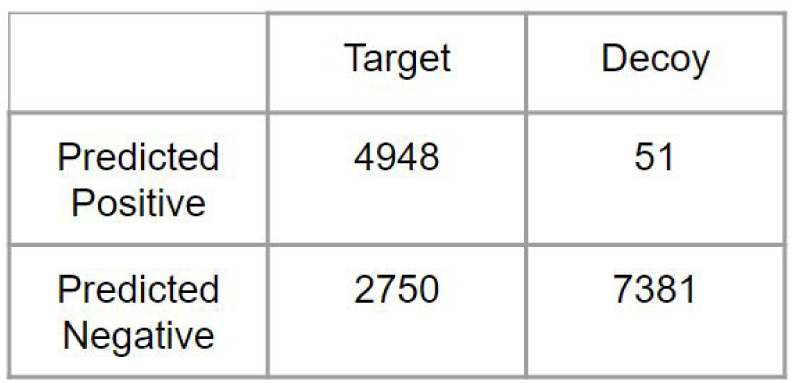
Confusion matrix of the QDA DIA extraction classifier.

|  | Target | Decoy |
| --- | --- | --- |
| Predicted Positive | 4948 | 51 |
| Predicted Negative | 2750 | 7381 |

### Spectral comparison among experimental DDA spectra, experimental DIA pseudo spectra, and predicted spectra from Prosit and LC-MS/MS property prediction tools

The iRT vs RT scatter plot of Figure 3 showed that the calibrated retention through the predicted iRT can be used effectively for DIA extraction. In supplemental figure S6, performance for iRT prediction for both Prosit(12) and LC-MS/MS property prediction(7) tools were compared.

We are now in a position to show in Table 2 that the predicted spectral library is suitable for extraction of chromatograms for the fragment ions. The DDA spectra referred to in Table 2 are the identified experimental spectra from DDA scans. The DIA spectra are pseudo spectra obtained by sum of all intensities in the DIA RT extraction peak windows. As expected, both predicted spectra from Prosit(12) or LCMSMS Property Prediction tools(7) match experimental DDA spectra well since both models were trained with DDA generated spectra. Although the 30% collision energy was specified in the Prosit prediction, the LC-MS/MS Property Prediction tools (without collision energy specification) matches to the DDA spectra better. LCMSMS Property Prediction tools correlates well with Prosit. The pseudo spectra from DIA extraction have a poorer correlation with the DDA spectra.

**Table 2.** Comparison between DDA, DIA experimental spectra and predicted spectra from Prosit and LCMSMS Property Prediction tools. Median Pearson Correlation Coefficient (median PCC) values are shown.

|  | DIA pseudo spectra | Prosit | LC-MS/MS<br>Prediction tools |
| --- | --- | --- | --- |
| DDA spectra | 0.893 | 0.978 | 0.987 |
| DIA pseudo spectra |  | 0.825 | 0.867 |
| Prosit |  |  | 0.981 |

### DIA Extraction of High Detectability Peptide Ions

The process to generate the high detectability spectral library is the same as that for the de Bruijn decoy spectral library, except this time the target human proteome sequences were used. Briefly, 551,628 peptides with detectability (computed by the AP3 program) greater than or equal to 0.6 were selected and their charge states were expanded by their survey charge state distributions. The high detectability spectral library of 704,609 peptide ions was constructed for interrogation of the DIA scans. The extracted features were classified using the QDA classifier trained with the DDA identified peptides and the downsampled high detectability de Bruijn decoy peptides.

Since the spectral libraries for both the DDA identified peptides and the high detectability peptides are generated from the same HCD spectral prediction tool and the same retention time calibration, the statistical behavior should be the same. Use of the same classifier and the same classification probability threshold to control the false discovery rate (1%) is justified. The total number of peptide ion identifications is 17,796, corresponding to 16,901 unique peptides, and 6,204 protein groups. When the whole de Bruijn decoy spectral library (577,487 entries) was used to extract DIA scans, 4,876 were declared as positive. The number of entries for the whole de Bruijn decoy spectral library per false positive hit (188 = 577,487/4,876) is smaller than that (152 = 7,781/51) for the downsampled decoy spectral library, indicating that the actual peptide FDR rate may be lower than 1.06%.

Although 16,901 of unique peptides identified here is a rather small number due to the factors discussed earlier, such suboptimal instrument method, inadequate extraction algorithms, and the high detectability selection algorithm, the number of 6,204 proteins inferred is respectable. Protein inference from peptide identification may be realized by classifying protein groups using peptide identification features, such as DeepPep(13), a deep learning based method. Here we can provide an upper bound of protein FDR of 2.4%, obtained by the following argument: 1.06% FDR for unique peptides results in 146 falsely identified peptide. Assuming all the 146 peptides represent 146 unique protein groups, the protein group FDR is 2.4% (=146/5673).

## DISCUSSION

Advancement in mass spectrometry instrumentation development continuously improves DDA and DIA experiment performance. As shown in Figure 1, the utilization of precursor RT-m/z space is poor even in the DIA portion. In the conventional ion isolation method, precursor ions of interest are isolated by a quadrupole (or ion trap) and other ions outside of isolation window are lost. The currently developed Online Parallel Accumulation–Serial Fragmentation (PASEF) method(14, 15), realized on a trapped ion mobility (TIMS) device allows for nearly 100% of precursor ion utilization. The high precursor utilization efficiency is achieved by storing ions in the TIMS device, effectively introducing a new ion mobility dimension in the RT-m/z space. With each splice of a given ion mobility, the isolation windows can occupy different regions. The projected final RT-m/z space can be filled up. Running in the DDIA mode, the PASEF method is expected to achieve unpreceded performance.

Neither MS instrument method nor data processing algorithms for theDDIA experiment presented here has been optimized. The product ions of the DDA scans should be detected with higher sensitivity and higher speed in an ion trap. The portion of time spent on DDA relative to that for DIA is a trade off between DDA and DIA performances. More DDA peptide identification gives a more accurate iRT vs RT calibration curve and a more accurate classifier for distinction between false and true DIA identifications. With a faster detection device, the DDA portion may be reduced. The DIA isolation window design was not optimal and a better design(5) should be implemented. The data processing algorithms are also primitive. The function for aggregation of extracted ion chromatograms with different power factors and their values were selected rather empirically without thorough investigation. In addition, the DIA extraction window width of ±3mins is quite wide, due to inaccuracy in iRT prediction. Nonetheless, we have demonstrated the usefulness for combining the two traditionally separated paradigms into a single LC-MS/MS experiment.

Interrogation of DIA scans with a large spectral library may result in more identification. However, the false discovery rate (FDR) control is more challenging. The machine learning field has developed many strategies to deal with the problem of “finding a needle in a haystack”, generally called anomaly detection. Deep learning based anomaly detection methods has been recently reviewed(16). In this work, random samples from the large decoy spectral library extraction should have reasonable statistical representation of that from the large decoy library. The False discovery rate (FDR) control with the de Bruijn decoy spectral library may also be highly conservative. We speculate that the high detectability decoy peptides were selected and many of them may have the same DIA behavior as real peptides: have the same precursor m/z (within the m/z tolerance) and the same fragment ions (also within the m/z tolerance). But the speculation needs to be rigorously investigated.

Traditionally, a large spectral library is prepared using (a) previously acquired experimental DDA spectra and (b) spiked-in iRT standard peptide for retention time calibration. As illustrated in Table 1, the predicted HCD spectra (from both Prosit(12) and LC-MS/MS property prediction tools(7)) are highly similar to the experimental DDA spectra. Although the iRT prediction accuracy is still far from optimal, it will be improved with more sophisticated deep learning models to incorporate different experimental conditions. The proof of concept DDIA experiment presented here suggests that it is possible to achieve high performance without the need for tedious spectral library preparation and experimental retention time calibration.

## Author contributions

S.G., M.M., and B.M. designed research; S.G. and P.T. performed research; S.G. and Z.H. contributed new reagents/analytic tools; S.G. analyzed data; and S.G. wrote the paper.

## ABBREVIATIONS

DDA: data dependent acquisition
DDIA: data dependent-independent acquisition
DIA: data independent acquisition
FDR: false discovery rate
iRT: indexed retention time
RT: retention time
QDA: quadratic discriminant analysis
ROC: receiver operating characteristic

## DATA AVAILABILITY

The code and data are provided in https://zenodo.org/deposit/3466997

## ACKNOWLEDGEMENT

* SG received partial funding for this work from the Pan-Canadian Proteomics Centre, a Genomics Technology Platform funded by Genome Canada. MFM was funded in part by Canada Research Chairs, Canada Foundation for Innovation, and National Sciences and Engineering Research Council of Canada. BM was funded in part by National Sciences and Engineering Research Council of Canada discovery grant and a GenomeCanada B/CB grant. SH, MFM, and BM receive funding from PCPC/Genome Canada.

## Notes

https://zenodo.org/record/3466997#.XaB2YyhKi70

