## supplemental_materials for "DDIA: data dependent-independent acquisition proteomics - DDA and DIA in a single LC-MS/MS run"

supplemental figure S1

### DDIA Data Processing 1

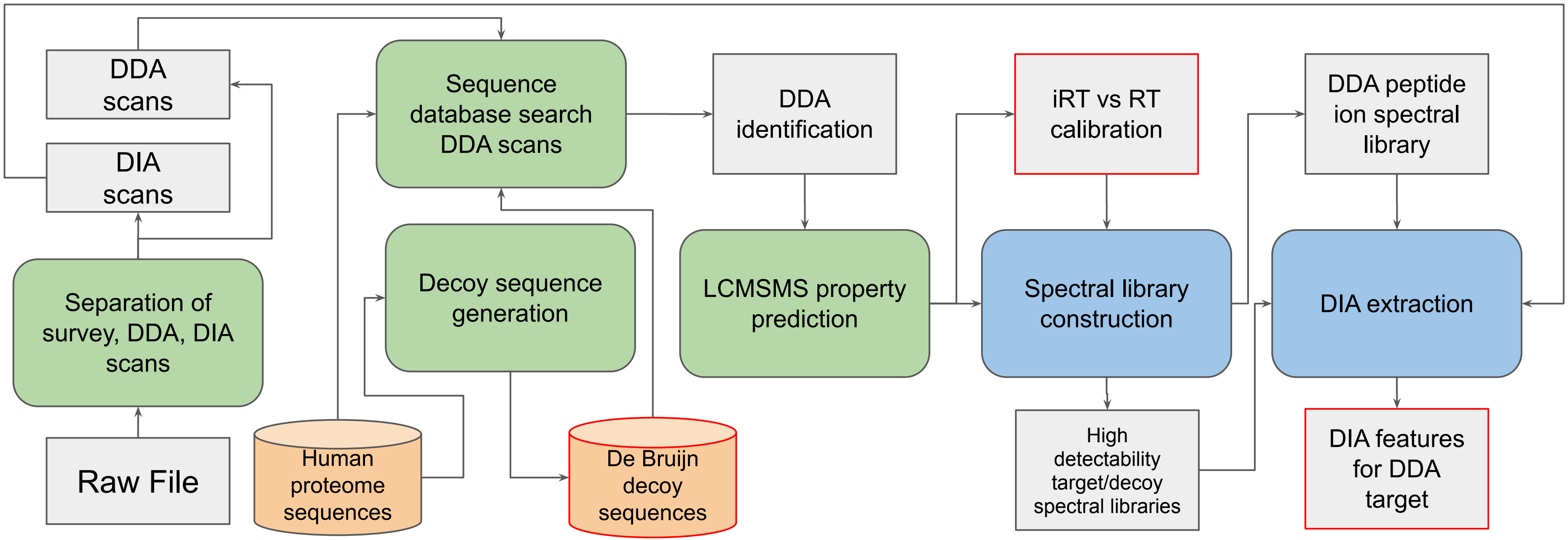


*Obtain necessary information from DDA data for DIA extraction*

Input: RAW file; Human proteome sequences

Output:

a. De Bruijn decoy protein sequences
b. iRT vs RT calibration curve
c. Target DIA features for training DIA classifier (DDA identified peptides)

supplemental figure S2

### DDIA Data Processing 2

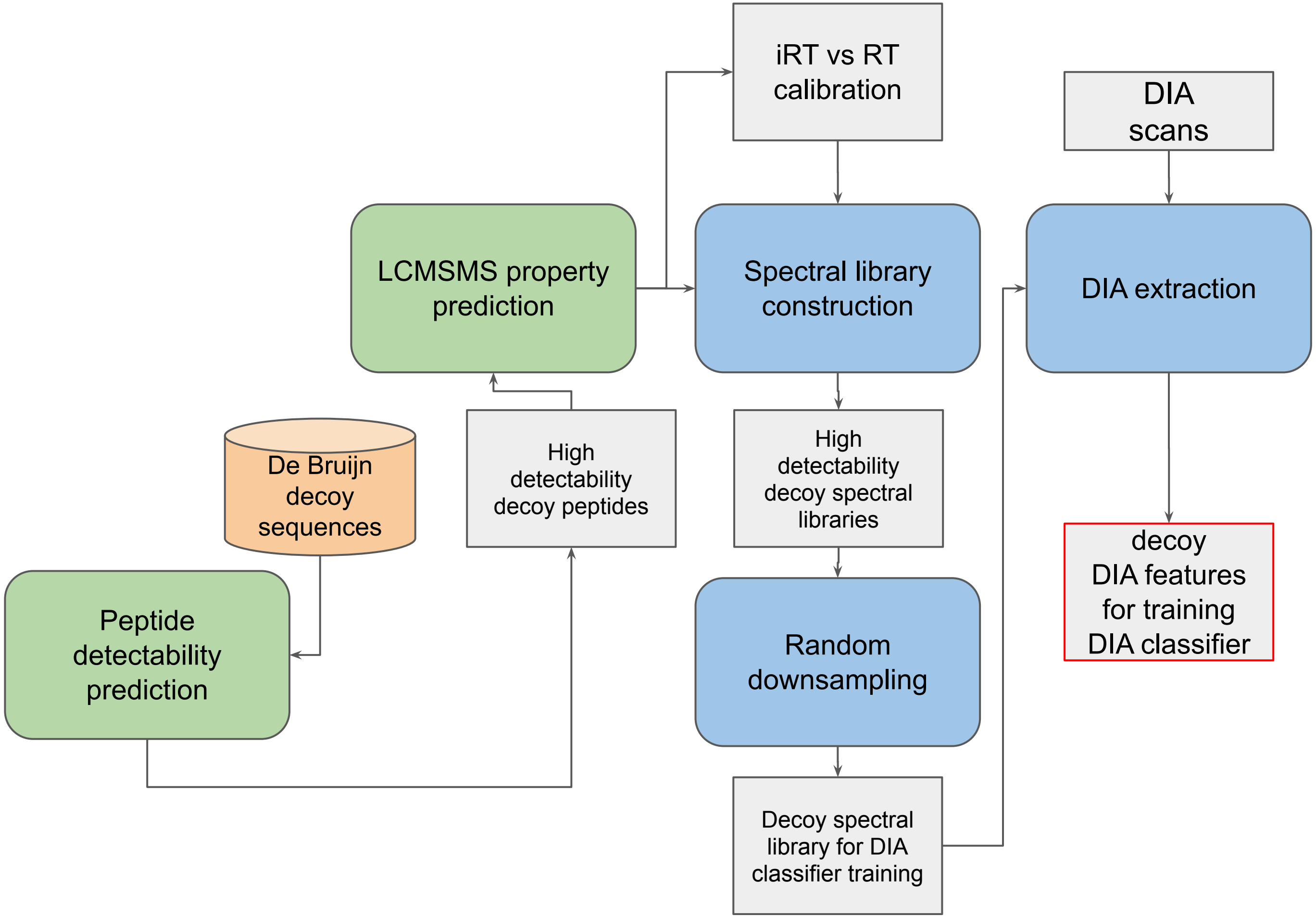


***<u>Obtain decoy DIA features for training DIA identification classifier</u>***

Input:

De Bruijn decoy protein sequences

Output:

Decoy DIA features

supplemental figure S3

### DDIA Data Processing 3

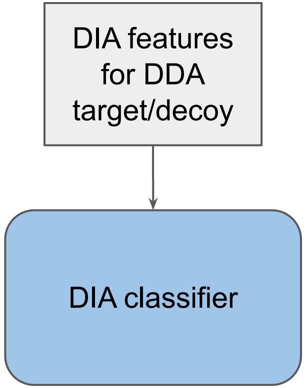


*Obtain DIA identification classifier*

Input:

Target DIA features from DDA identified peptide ions and decoy DIA features from downsampled high detectability peptides of de Bruijn decoy sequences

Output:

DIA identification classifier and FDR threshold

DDA target (7781)/DBD downsampled decoy(7781)

Class probability threshold = 0.996

- TP = 3739/1209 (training/testing)
- FP = 38/13
- TN = 5520/1861
- FN = 2050/700

FDR rate = 1.01%/1.06%
ACC = 0.816/0.812

supplemental figure S4

### DDIA Data Processing 4

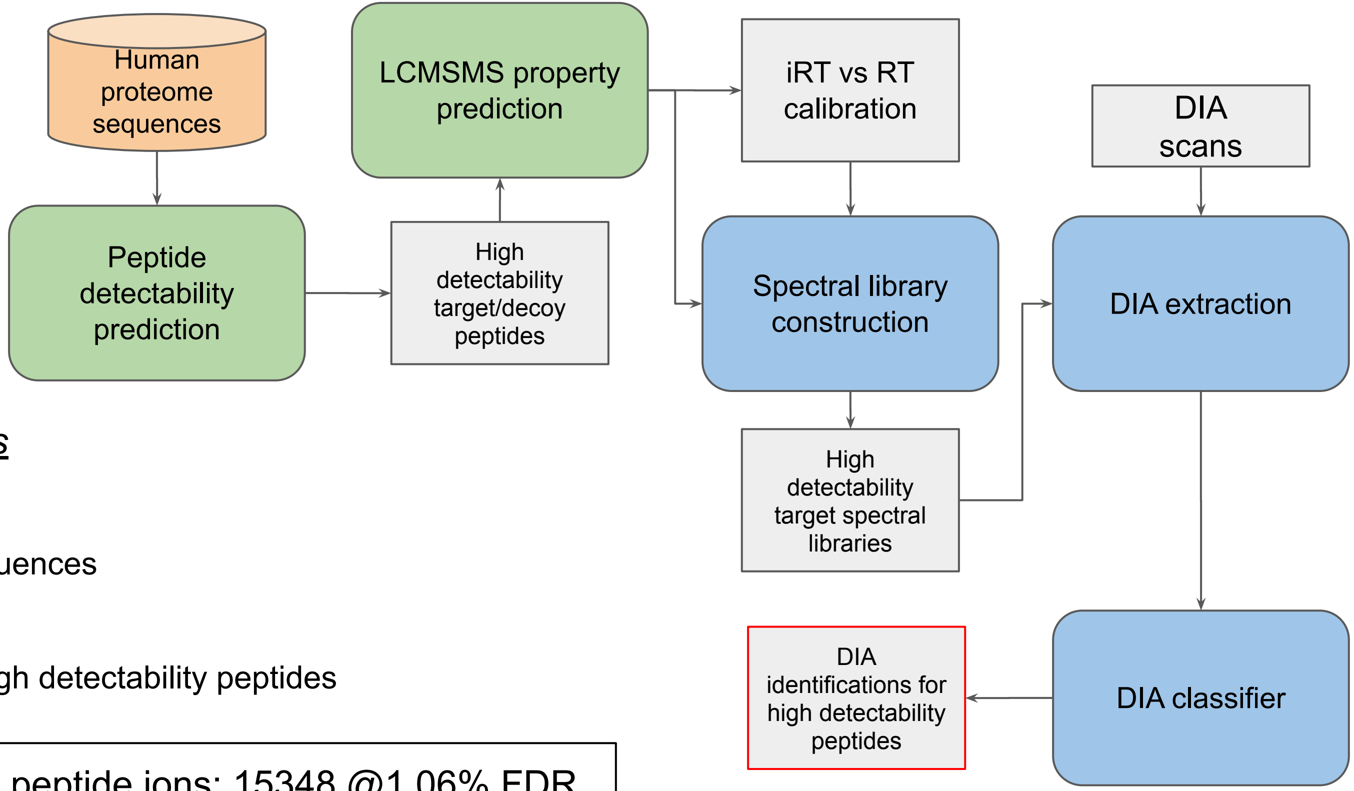


*__Obtain DIA identifications__*

Input:

Human proteome sequences

Output:

DIA identification of high detectability peptides

High detectability peptide ions: 15348 @1.06% FDR
Unique peptides: 14572
Protein groups: 5673, FDR<2.6%

supplemental figure S5

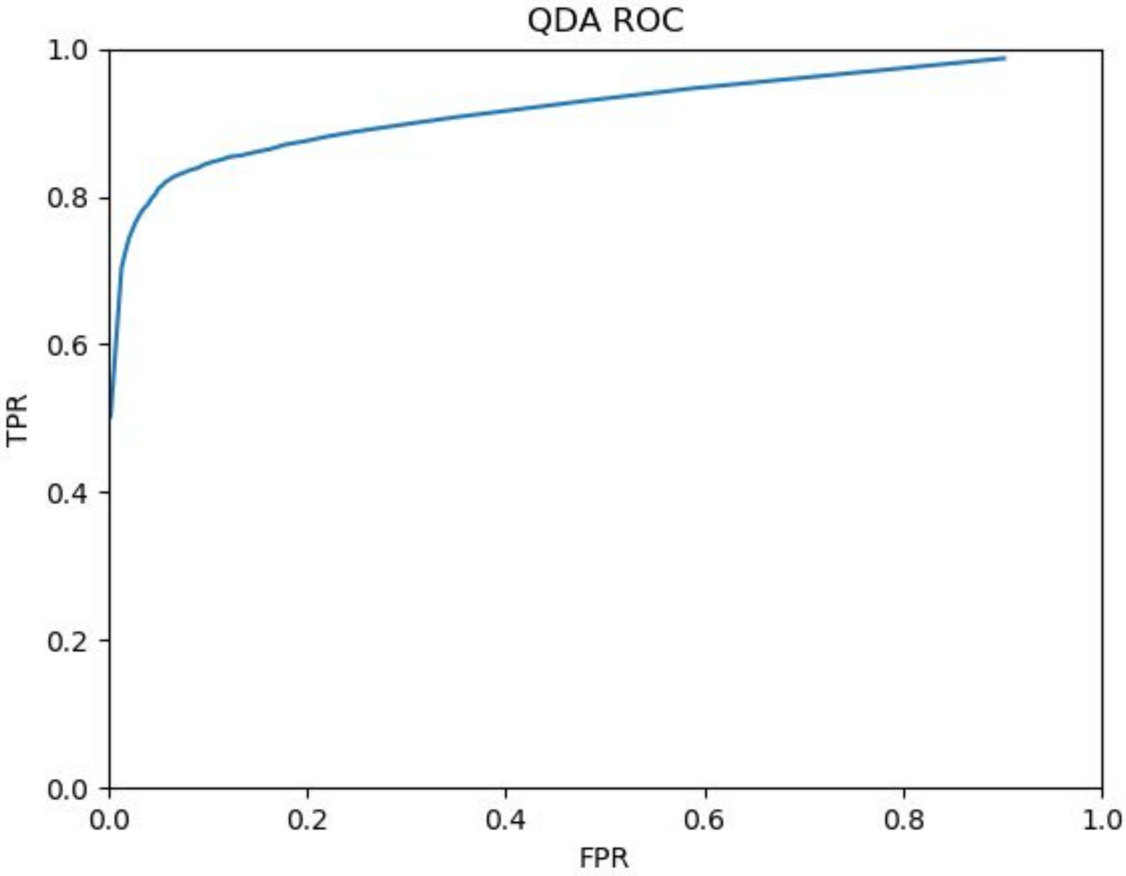

supplemental figure S6

### Predicted iRT vs RT Calibration

Prosit (ProteomeTools)

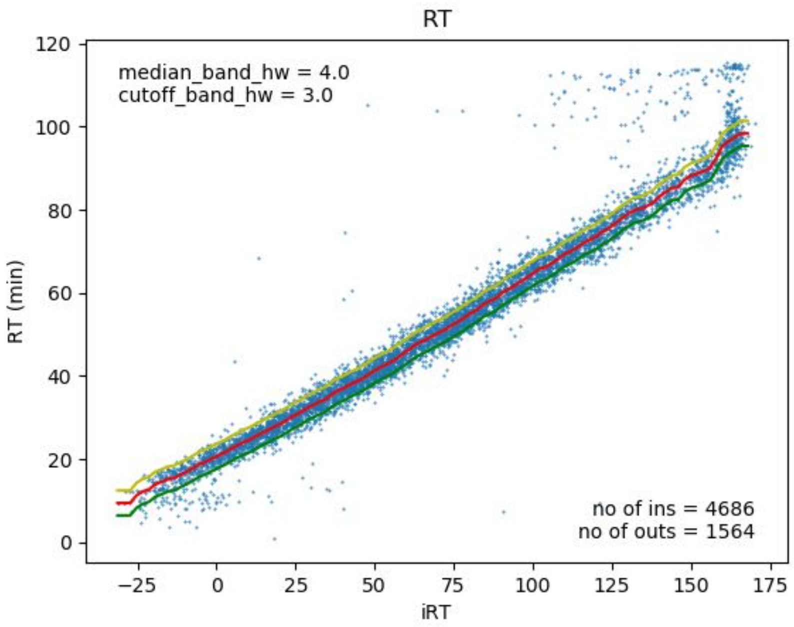


NatMethods2019v16p519-525

Model 10

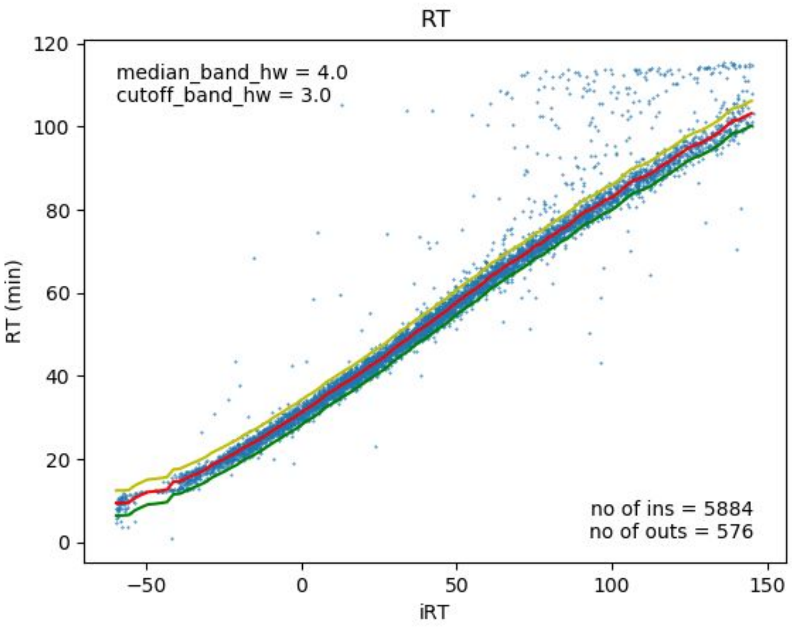


MolCellProteomics2019Jun 27

DIA Extraction

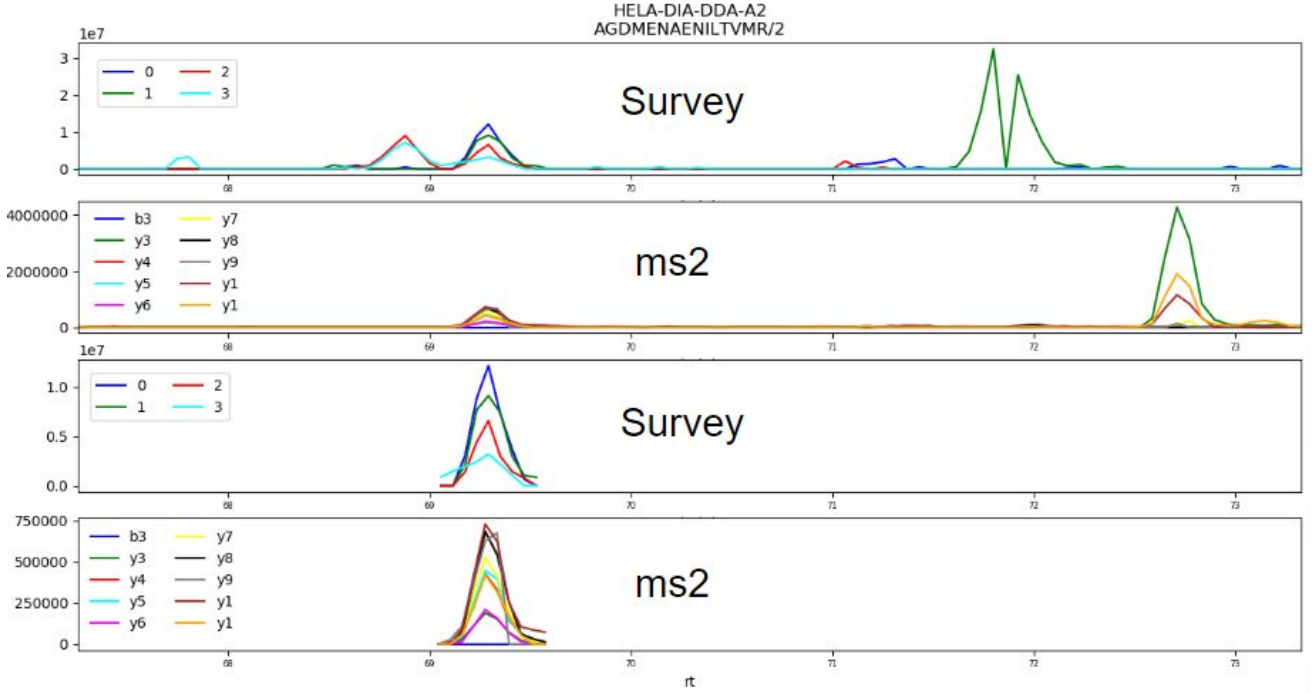


Supplemental figure S7. Ion chromatograms of AGDMENAENILTVMR/2 ion. Panel 1 is the isotope ion chromatograms for the peptide precursor ions in the ±3 mins RT extraction window. Panel 2 is the fragment or product ion chromatograms in the full RT extraction window. Panels 3 and 4 are ms and m2 chromatograms after peak detection. DIA extraction features were calculated from the truncated chromatograms.

Survey XICs were aggregated into (a) theoretical isotope distribution projected chromatogram (dot product of experimental values with the theoretical isotope distribution)(Wang et al. 2015) and (b) theoretical isotope distribution pairwise distance chromatogram (one minus the cos distance between experimental values and the theoretical isotope distribution). The pairwise distance in this definition has a mimal value of 0 (least similar) and maximal value of 1 (most similar).

In a similar fashion, ms2 or DIA XICs were also aggregated into projected and pairwise distance chromatograms. Instead of using the theoretical isotope distributions as a template in the survey extraction, the spectral library HCD spectra was used.

The four aggregated chromatograms were further aggregated into a single chromatogram by multiplication together with different powers of each: 1 for ms2 pairwise distance, 0.5 for survey pairwise distance, 0.1 for ms2 dot-product, and 0.01 for survey dot-product. The power coefficients were selected rather empirically: the higher the value the more weight is given. Survey intensities are generally orders of magnitude higher than those for ms2. Those power "weights" were rather superficially selected. But their optimal values may be learned with a machine learning method.

Wang, J., Tucholska, M., Knight, J. D. R., Lambert, J.-P., Tate, S., Larsen, B., Gingras, A.-C., and Bandeira, N. (2015) MSPLIT-DIA: sensitive peptide identification for data-independent acquisition. *Nat. Methods* 12, 1106–1108

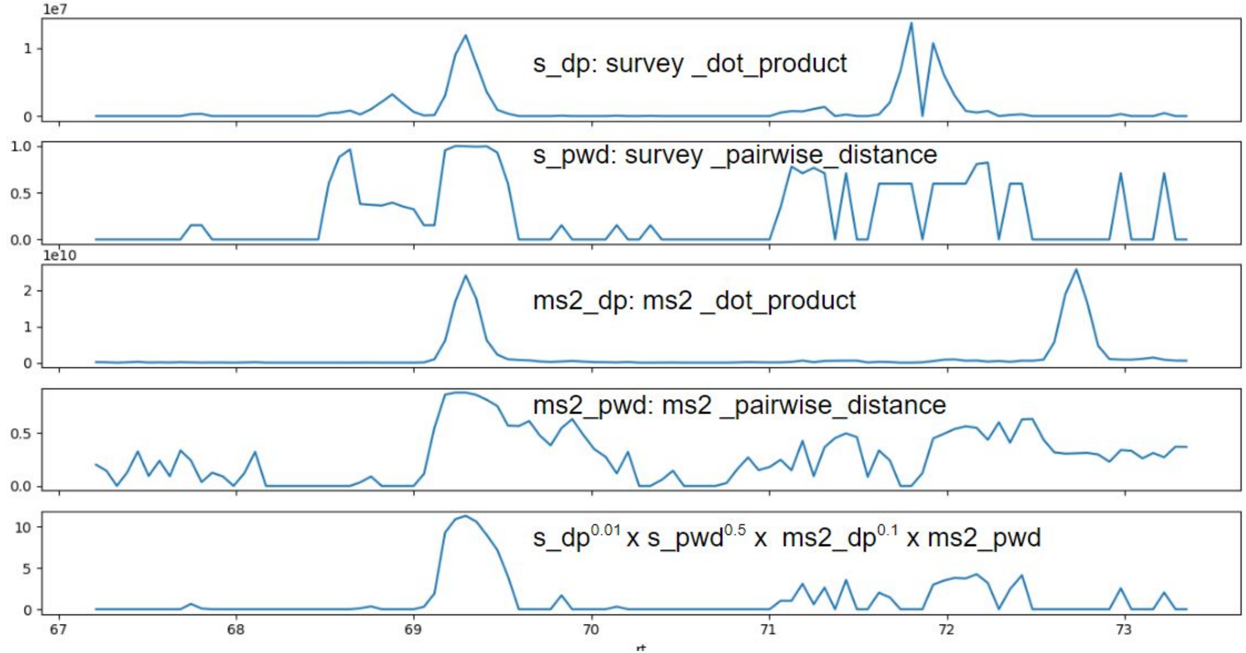


Supplemental figure S8. aggregated ion chromatograms. Panel 1: constructed from the dot product between the experimental survey data and theoretical isotope distribution. Panel 2: one minus cos pairwise distance between the experimental survey data and theoretical isotope distribution. Panel 3: the dot product between the experimental ms2 data and the spectral library product ion intensities. Panel 4: one minus cos pairwise distance between the experimental ms2 data and the spectral library product ion intensities. Panel 5. Aggregated ion chromatogram from Panels 1-4, by multiplication with different power "weights".

The final aggregated chromatogram (called peptide ion chromatogram) was smoothed and 20% intensity threshold applied.  The highest peak was chosen to compute the following six features: averaged transformed survey dot product, averaged survey pairwise distance, averaged transformed ms2 dot product, averaged ms2 pairwise distance, average pairwise distance between survey and ms2, retention time difference between survey and ms2.

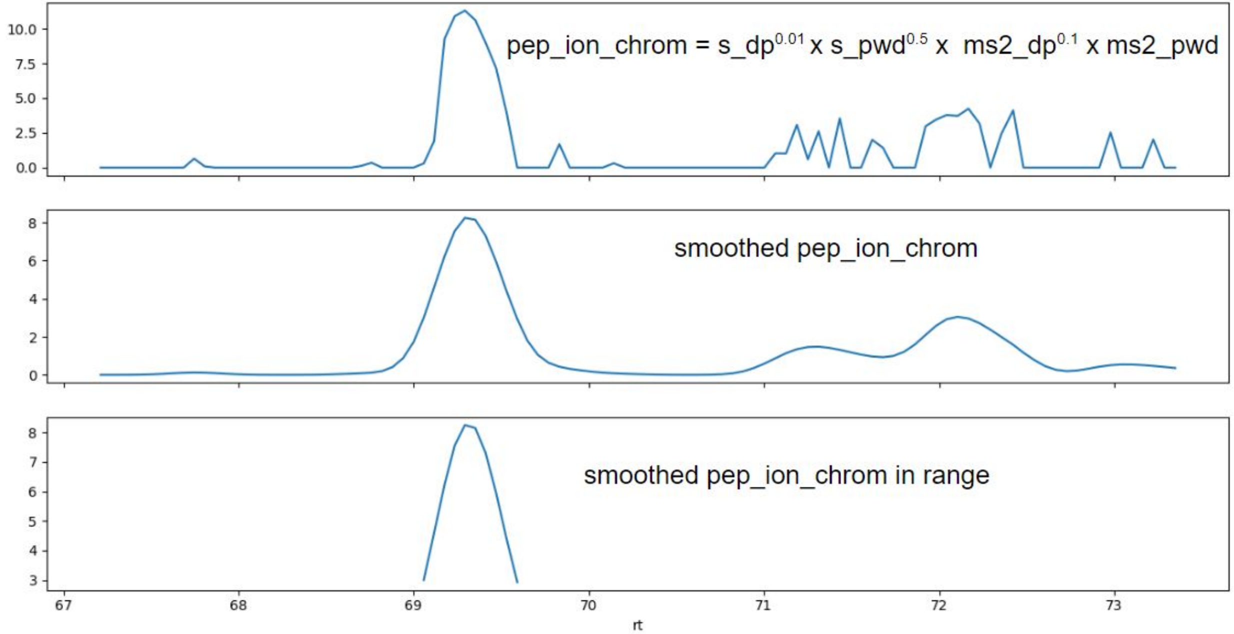


Supplemental figure S9.  Smoothing of peptide ion chromatogram.  Panel 1: the aggregated peptide ion chromatogram (the same as the last panel in supplemental figure S8).  Panel 2: smoothed chromatogram of Panel 1.  Panel 3: range-selected or peak detected or truncated peptide ion chromatogram.  In supplemental figure S7, the lower two panels were produced after truncation using Panel 3 of this figure.
